# Universality of spatial livestock disease transmission patterns

**DOI:** 10.1101/2022.07.14.500003

**Authors:** Gert Jan Boender, Thomas J. Hagenaars

## Abstract

The risk of epidemic spread of diseases in livestock poses a threat to animal and often also human health. Important for the assessment of the effect of control measures is a statistical model quantification of between-farm transmission during epidemics. In particular, quantification of the between-farm transmission kernel has proven its importance for a range of different diseases in livestock. In this paper we explore if a comparison of the different transmission kernels yields further insight. Our comparison identifies universal features that connect across the different pathogen-host combinations and thereby provide generic insights.

Comparison of the shape of the spatial transmission kernel suggests that, in absence of animal movement bans, the distance dependence of transmission has a universal shape analogous to Lévy-walk model descriptions of human movement patterns. Also, our analysis suggests that interventions such as movement bans and zoning, through their impact on these movement patterns, change the shape of the kernel in a universal fashion. We discuss how the generic insights obtained can be of practical use for assessing risks of spread and optimizing control measures, in particular when outbreak data is scarce.

## Introduction

Epidemics of highly contagious diseases in livestock such as Foot and-Mouth Disease (FMD) and high pathogenic avian influenza (HPAI) can have tremendous socio-economic consequences as well as devastating effects on animal health ^1,2^. For developing effective contingency planning for the control of such epidemics, it is essential to use the data from previous epidemics to gain as much insight as possible in the quantitative characteristics of transmission, in particular of between-farm transmission ^3^. These characteristics may include the observed effect of control measures such as animal movement standstill or zoning ^4^. By means of using mathematical models fitted to describe these characteristics, past or current outbreaks in a given country are often being used to make extrapolations to current or future transmission risks in that same country ^5,6^. For countries with no earlier or no informative outbreak, one may resort to extrapolation from epidemic patterns observed in other countries, addressing uncertainties about model representativity for the country of interest e.g. by exploring the sensitivity of the model outcomes to possible differences in parameter values ^7^. For emerging diseases for which there is no previous epidemic that can be analysed, what can we learn from the patterns observed in epidemic data of other livestock diseases? Here we present a general framework for the spatial transmission of livestock diseases that can help to underpin model extrapolations between control strategies, between countries, and also between diseases. It is built on the comparison of epidemic transmission risk patterns across a range of livestock diseases.

Our framework uses the transmission kernel as a central element in the modelling approach. Transmission kernels describe the distance-dependent probability of transmission from an infected to a susceptible farm, and have been used to describe the between farm transmission of different animal diseases ^5,6^. Use of a transmission kernel avoids the modelling of specific transmission pathways where these are poorly known, and allows both the construction of risk maps as well as model simulation studies of the effectiveness of control measures ^5,6,8,9^. The transmission kernel also facilitates the comparison of the distance-dependent characteristics of transmission between epidemics, between diseases, and between phases in one and the same epidemic differing in applied control measures ^10-12^.

For the interpretation of the comparative kernel analyses results we can build on a body of literature that uses spatial kernels to describe movement and dispersal patterns ^13,14^. One of the elements from this literature is a distinction between thin-tailed (i.e. exponentially bounded) and fat-tailed (i.e. power-law) kernels ^15^. Whereas thin-tailed kernels generate ‘diffusive’ dispersal patterns with constant-speed travelling waves, a fat-tailed kernel produces ‘super-diffusive’ behavior lacking a finite velocity and yielding a patchy dispersal pattern ^15-18^. In the description of animal movement, thin-tailed kernels are a signature for an underlying Brownian random walk and fat-tailed kernels for Lévy-walk patterns; based on this distinction many studies demonstrate Lévy-walk patterns for animal movement ^19^. Also human mobility appears to follow a Lévy walk ^20,21^.

An important subtlety described by the so-called ‘truncated’ Lévy-walk model is the phenomenon that the fat-tailed/power-law behaviour is truncated by an exponential decline setting in above a cut-off distance scale. This truncated version of the Lévy-walk model describes super-diffusive movement on a distance small compared to, and diffusive movement on a scale large compared to, the cut-off distance ^22^. In the case of human mobility, such a cut-off distance scale may reflect the existence of an area within which most of the movements are confined, e.g. an urban area for the movements of commuters ^23^. For human travel mobility patterns the power-law exponent in the truncated Lévy walk appears to have a universal value of about 1.6 ^17,20^.

The aim of this article is to yield insight into the similarities and differences between a range of spatial patterns of animal disease transmission by comparison of the corresponding transmission kernels. We will use one and the same kernel parametrisation throughout such that parameter values can be directly compared. In our previous kernel analyses of between-farm transmission we have mostly adopted a so-called Cauchy form to parameterize the kernel, motivated in part by the fact that it was identified as having the lowest AIC amongst a set of alternatives studied in ^6^. Recent analyses however suggest that the (non-truncated) Lévy-walk kernel produces an even better fit to spatial transmission data from certain livestock disease epidemics ^10,24^.

Therefore we re-analysed eight datasets available to us using the Lévy-walk kernel. In addition to this, we assembled from the literature three Lévy-walk kernels estimated for other animal disease epidemics. The resulting set of eleven Lévy-walk kernels allows us to compare the results to see if similarities extend across a broad set of livestock disease transmission patterns. A descriptive overview of the epidemics included is given as part of the Materials and Methods. In the Results, the estimated Lévy-walk model parameters for each epidemic are presented and correlated with applied control strategies. In the Discussion, we present the insights obtained and their practical relevance for assessing risks of spread and optimizing control measures, in particular when outbreak data is scarce.

## Material and Methods

To investigate to which extent observed between-farm transmission patterns can be categorized and interpreted using insights from the study of movement patterns, we assemble and compare the Lévy-walk kernel fits to 11 different between-farm transmission data sets. These datasets are listed in Table 1. Three of the datasets were already analyzed using Lévy-walk kernels in the literature, namely two parts of the UK 2001 FMD epidemics and the 2010 FMD epidemic in Japan ^10,24^. The remaining eight datasets of livestock epidemics or parts of epidemics ^6,8,11,12,25,26^ are reanalyzed here by fitting truncated Lévy-walk kernels. These eight comprise the 2006 Q fever epidemics in the Netherlands, two parts of the Swine Vesicular Disease (SVD) epidemics in 2006/2007 in Italy, the 2001 FMD epidemic in the Netherlands, the 2003 HPAI epidemic in the Netherlands, the 1997/1998 Classical Swine Fever (CSF) epidemic in the Netherlands, and the 2006 Blue Tongue (BT) epidemics in Germany and Belgium. These eight epidemic datasets were previously analyzed using the Cauchy form of the kernel

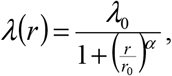

**Table 1.** List of 11 epidemic data sets analyzed, ordered by applied control strategy (in order of increasingly stringent measures) and year.

| Nr | Disease | Year | Country | Strategy | Kernel model of existing analysis |
| --- | --- | --- | --- | --- | --- |
| 1 | FMD<sup>10</sup> | Before 23<sup>rd</sup><br>February 2001 | UK | NMNZ | Lévy Walk |
| 2 | SVD<sup>12</sup> | 2006 | Italy | NMNZ | reference |
| 3 | Q fever<sup>25</sup> | 2007-2010 | The Netherlands | NMNZ | reference |
| 4 | BT<sup>11,24</sup> | 2006 | Belgium | NMZ | reference |
| 5 | BT<sup>11</sup> | 2006 | Germany | NMZ | reference |
| 6 | CSF<sup>26</sup> | 1997-1998 | Netherlands | M | reference |
| 7 | FMD<sup>10</sup> | Post 23<sup>rd</sup><br>February 2001 | UK | M | Lévy Walk |
| 8 | FMD<sup>8</sup> | 2001 | The Netherlands | M | reference |
| 9 | HPAI<sup>6</sup> | 2003 | The Netherlands | M | reference |
| 10 | SVD<sup>12</sup> | 2007 | Italy | M | reference |
| 11 | FMD<sup>24</sup> | 2010 | Japan | M | Lévy Walk |

in which λ^_0_^ represents the amplitude of the transmission kernel (defined as the transmission hazard for small distances), r_0_ a characteristic distance, and α the long-distance scaling exponent. In this paper we keep the Cauchy kernel results for reference, to compare the fits to those using the Lévy-walk kernel. We use the following parametrization for the truncated Lévy-walk kernel ^20^:

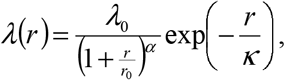

in which the additional parameter κ is a cutoff distance truncating the Lévy-walk behavior. The parameter estimates are obtained Maximum Likelihood (ML) estimation, and confidence bounds using the likelihood-ratio test. As the kernel amplitude λ_0_ does not inform about the distance dependence of transmission, we only report its estimated values in the Supplementary Information for completeness. We note that the estimate for the cutoff distance κ is only informative if it is smaller than the ‘extent’ of the area spanned by the dataset; if larger, then no truncation is detected on the distance scales covered by the data.

During the 11 epidemics the control strategy was either a movement ban (M), or no movement ban but zoning (NMZ), or no movement ban nor zoning (NMNZ). Zoning was applied in the 2006 BT epidemic in Germany with a typical zone radius of about 20 km and in the 2006 BT epidemic in Belgium, with spillover to the Netherlands and France, in which during the main part of the epidemics Belgium was considered to be one single zone corresponding approximately to a radius of about 140 km (half the extent of the country) ^11,27^. Using the Akaike’s Information Criterion (AIC) the model fit of the Lévy-walk kernel was compared that of the reference (i.e. Cauchy) kernel (AIC_0_). Differences in AIC’s are considered to be significant if larger than 2 ^28,29^.

In order to objectively identify categories of epidemic transmission patterns we carried out a hierarchical clustering analysis using the estimated parameter pairs {α, κ} ^30^. Here we used a Euclidian distance function after rescaling both parameters to take values between 0 and 1. The rescaling was carried out using a logistic scaling function, in which for each of the two parameters the scaling factor was set to the median value across all ten estimates. The p-value of the categorization into clusters is calculated by basic combinatorics.

## Results

The estimated parameters for the Lévy-walk kernel for the 11 epidemics are given in Table 2. Amongst the eight epidemics we re-analyzed here, only epidemics 4 and 5 correspond to a finite spatial limitation κ. For epidemics 2, 3, 6 and 8-10, the estimation yielded values for the cutoff parameter κ that were much larger than the spatial extent of the datasets and that were poorly converging, indicating that super-diffusive behavior extends across all length scales in the dataset. Therefore, for these epidemics, in the final ML estimation procedure we set the factor 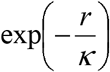 equal to 1 (corresponding to assuming κ = ∞) to remove the convergence problems. For the epidemics 1-5 (i.e. the epidemics without movement ban) the estimate of the scaling exponent α ranges between 0.66 to 1.83 (with the majority of values between 1.45 and 1.83) and for the epidemics 6-11 (with movement ban) it ranges between 2.36 and 2.68. The AIC and the AIC_0_ are significantly different (difference>2) only for epidemics 4 and 5 (truncated Lévy-walk kernel performing better than the reference kernel). For the kernel estimation results taken from the literature (epidemics 1, 7 and 11) that literature gives results for non-truncated Lévy-flight kernels; i.e. no estimates for κ are available.

**Table 2.** Estimations for the 11 epidemics of the Lévy-walk kernel parameters *r*_0_, *α* and *κ* (with confidence bounds between brackets), the corresponding AIC, the AIC_0_ for the reference kernel estimation, and the applied control strategy. For the kernel parameters of the transmission kernel for FMD in Japan (epidemic 11) no confidence bounds were reported in ^24^.

| Nr | Strategy | $r_0$ (km) | $\alpha$ | $\kappa$ (km) | AIC | AIC<sub>0</sub> |
| --- | --- | --- | --- | --- | --- | --- |
| 1 | NMNZ | N/A | 1.72 (1.54,1.93) | N/A | N/A | N/A |
| 2 | NMNZ | 0.36 (0.02,1.64) | 1.83 (1.45,2.39) | ∞ | 437.5 | 436.632 |
| 3 | NMNZ | 2.5 (0.48,11.1) | 1.45 (1.1,2.1) | ∞ | 699.47 | 698.566 |
| 4 | NMZ | 1.32 (0.28,3.44) | 1.70 (1.39,2.09) | 173 (90,932) | 24125.3 | 24130.8 |
| 5 | NMZ | 0.18 (0,18.7) | 0.66 (0.38,2.13) | 25.3 (20.3,45.9) | 16767.9 | 16795.5 |
| 6 | M | 0.64 (0.43,0.96) | 2.56 (2.36,2.82) | ∞ | 6800.09 | 6801.96 |
| 7 | M | N/A | 2.68 (2.58,2.78) | N/A | N/A | N/A |
| 8 | M | 0.76 (0.17,2.64) | 2.36 (1.86,3.21) | ∞ | 505.884 | 505.015 |
| 9 | M | 2.7 (1.2,6.1) | 2.53 (2.03,3.46) | ∞ | 3089.80 | 3090.99 |
| 10 | M | 0.78 (0.003,4.05) | 2.47 (1.75,3.98) | ∞ | 206.852 | 205.017 |
| 11 | M | 0.58 | 2.47 | N/A | 2093.08 | 2094.32 |

In Fig 1 we show the dendrogram result the hierarchical clustering of the value pairs {α, κ} for the all 11 epidemics. This dendrogram indicates 2 clusters in which cluster 1 could subdivided in 2 sub-clusters. The clusters coincide completely with the arrangement according to strategy (P<0.001): The epidemics 1-3 with strategy NMNZ are all located in cluster 1a, the epidemics 6-11 with strategy M are all located in cluster 1b, and the epidemics 4 and 5 with strategy NMZ are all located in cluster 2.

**Figure 1.**
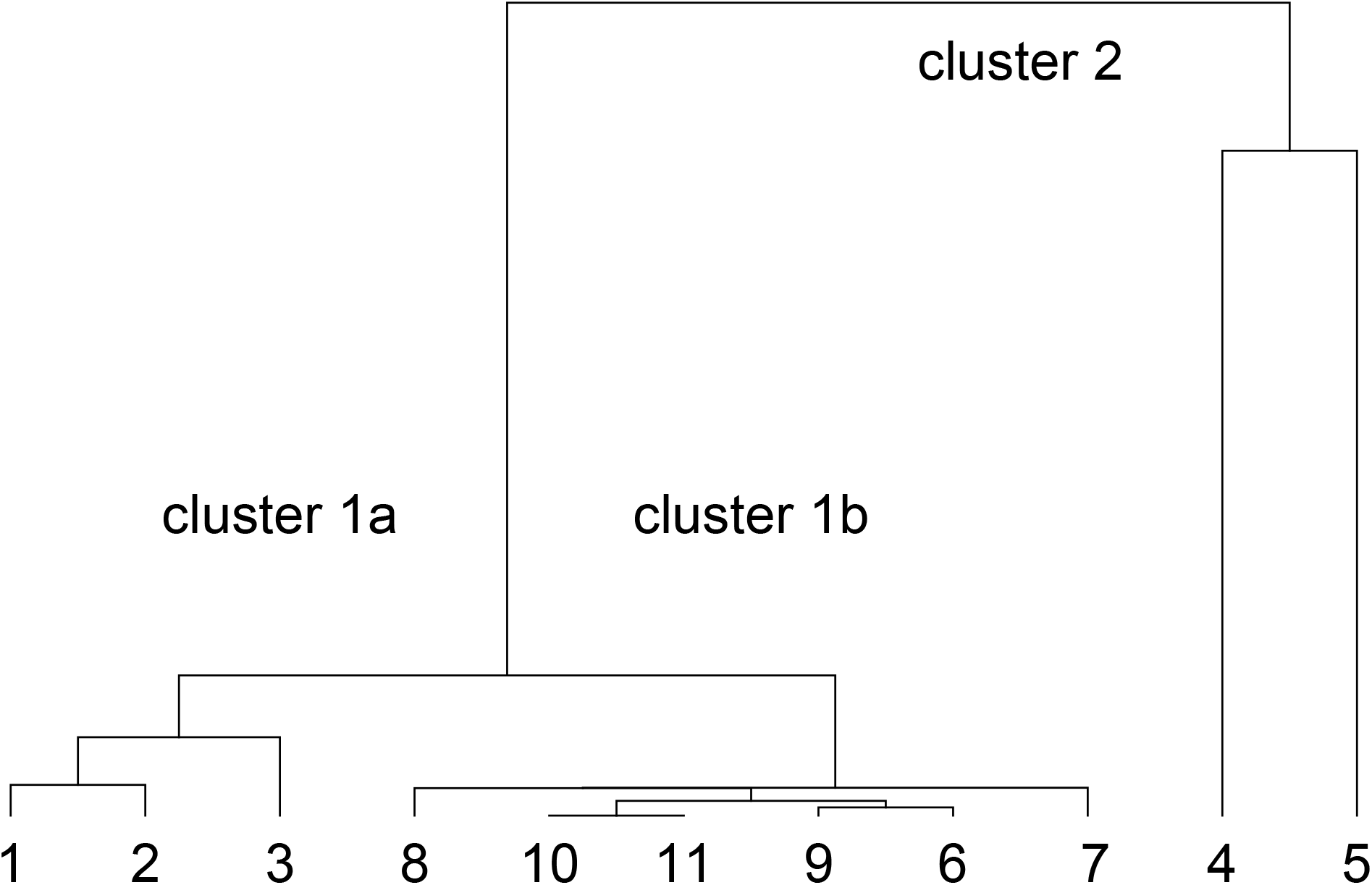
Dendrogram of the hierarchical clustering of the value pairs {*α, K*} for the 11 epidemics; two main clusters 1 and 2 are indicated, and within cluster 1 the two (sub)clusters 1a and 1b could be distinguished.

## Discussion

We found a perfect separation of the ranges of estimates for the scaling exponent α between strategies with and without movement ban (Table 2). We observe that for the strategies without movement bans the scaling exponent is most often close to (and never significantly different from) the universal value of approximately 1.6 identified in analyses of human travel mobility patterns. This suggests that in absence of movement bans, the distance dependence of between-farm transmission is strongly determined by the pattern of between-farm animal transports; and that this pattern is characterized by a scaling exponent taking a value close to the universal value for human mobility. Indeed, modelling suggests that the power-law scaling of movements of infected individuals would be directly reflected in the spatial transmission pattern ^31^. In line with this interpretation, we observe that only for the two epidemics with zoning measure (NMZ), the inclusion of the cutoff parameter *κ* leads to a better fitting model, suggesting that the zoning measure restricts the universal power-law dependence to distances in the order of the size of the zone. The applied protection zone of the 2006 BT epidemic in Germany (epidemic 11) was about 20 km and if zones overlapped they were merged to one larger zone ^11^. The estimate for the cutoff distance *κ* epidemic 5 is 25.3 (20.3,45.9) km which we deem fully consistent with these zoning measures. The estimate for the cutoff distance in the 2006 BT epidemic in Belgium (epidemic 4) is 173 (90,932) km; as in this analysis (in contrast to the previous analysis reported in ^11^) the populations of the Netherlands and France were also included, this 173 km represents an informative distance. This estimate is consistent with Belgium being declared one protection zone with the ‘radius’ of the country being about 140 km. The close correspondence, in both cases 4 and 5, between zoning measure and kernel shape further underpins the interpretation that constraining animal transports to within defined zones limits the dominant transmission route to distances within the extent of the zone. We note that although the analyses of the epidemics 1-3 and 6-11 do no produce a finite cutoff parameter, this is still consistent with the expectation that imposed export bans did produce a zoning effect on the scale of the country in question, as such an effect is not identifiable in an analysis including only the farm population within that country.

We observe that for the six epidemics with movement bans the estimated scaling exponent ranges between 2.36 and 2.68 which is clearly different from the range (0.66-1.83) estimated for the 5 epidemics without movement ban, the difference being about 1.0. The fact that the imposition of a movement ban produces such a dramatic shift of α, implies that in absence of a movement ban, the longer-distance disease transmission is mainly driven by animal transports. Other possible transmission routes, such as human and fomite movements unrelated to animal transports, movements of wild animals and wind-borne virus dispersal, together may only play a minor role in the longer-distance disease transmission as long as no movement ban is in place. After imposition of a movement ban these other possible transmission routes are the ones remaining, and are yielding a scaling exponent that, compared to the situation without movement ban, seems to be increased by an amount of about 1.0. This difference of about 1.0 can be interpreted if we make two assumptions. First, we assume that the dispersal or movements underlying the remaining transmission routes are occurring predominantly in random directions, i.e. not necessarily directed towards a neighboring farm. Second, we assume that these dispersal or movement processes are also described by a scaling exponent of approximately 1.6. The difference of about 1.0 can then be interpreted as a 1/r dilution factor that (in two dimensions) arises from the condition, in order for transmission to be possible, that the random direction of dispersal or movement of infectious material matches the direction towards specific (farm) locations; this in contrast to transmission via animal transports that are by definition directed towards a given (farm) destination ^31^. Regarding the first assumption, we note that the indirect between-farm contacts occurring when e.g. feed delivery or egg collection trucks visit multiple farms on the same day, are clearly directed to the farms as destinations. I.e. these types of contacts do not correspond to movement with a random direction. This means that our assumption corresponds to a situation where only a minor part of infections can be attributed to these types of contacts, as was the case for epidemic 9 of Table 1 ^32^. Regarding the second assumption, transmission via human movement, e.g. the route of a passerby accidentally connecting two farms, fits well into this interpretation as, Lévy-walk patterns with a scaling exponent of 1.6 have been identified for human movement ^20^. This interpretation would thus support the hypothesis that ‘random’ human movement is the most important between-farm transmission route once animal movement bans are in place. However, the value of 1.6 does not exclusively point towards human movement. Concerning movement of wild animals, it is known of some species that they also move according a Lévy-walk pattern with a scaling exponent of about 1.6 ^33^. Concerning wind, specific atmospheric conditions can lead to aerial dispersal following a Lévy-walk pattern with a scaling exponent of about 1.5 ^34^. In addition, we note that our interpretation also implicitly assumes that the movement of infectious material between farms is fast enough such that pathogen survival is not influencing the scaling of transmission with distance.

Our results show that after imposing a movement ban, transmission retains its super-diffusive character, although the movement ban does reduce the longer-distance transmission risks by increasing the scaling exponent α from around 1.6 to around 2.6. As long as transmission has a super-diffusive character (power-law dependence on distance), it will difficult to spatially confine it using local control measures such as ring culling of ring vaccination.

The analyses in this paper provide a conceptual framework for analysis of spatial livestock disease spread and control. In addition to its relevance for interpreting past epidemic patterns and underpinning model extrapolations from these patterns, our approach provides a framework for assessing the spatial transmission of emerging livestock diseases, i.e. diseases for which no previous epidemic is available to estimate kernel parameters. As a prior model, the between-farm transmission could be assumed to follow a Lévy-walk kernel with parameter values as follows. The exponential power α would be assumed to be about 2.6 if a movement ban is put in place and 1.6 without a movement ban. The spatial limitation κ would be assumed to equal the radius of the protection zone in case no movement ban is implemented and the radius of (a circular area approximating) the country when import bans are imposed by neighboring countries. According to Table 2, the scaling distance r_0_ seems to be of the order of 1 km for all epidemics included from Western European countries. The remaining information needed for the Lévy-walk kernel, namely the value of the (relative) amplitude of the between-farm transmission (λ_0_) for the emerging disease in question, could be treated as a scenario parameter, to be varied within a plausible range. The resulting prior model can be used for analyses that can comprise risk maps for the spread of the emerging disease as well as more detailed evaluation of control measures by means of kernel model simulations ^6,9^.

## Data availability

All data generated and analyzed during this study are included in this published article and its Natuementary Information file.

## Author contributions

Both authors wrote and reviewed the main manuscript. GJB performed the data analysis and visualized results in the figure. Both authors contributed to the interpretation and conceptualization.

