## Supplementary Table S1 for "Universality of spatial livestock disease transmission patterns"

**Table S1** The re-estimated Lévy-walk kernel parameter $\lambda_0$ (with confidence bounds between brackets) for the 9 epidemics in Table 1 with kernel model of existing analysis 'reference'.

| Nr | $\lambda_0$ |
|---|---|
| 2 | 0.0066 (0.0013,0.3) |
| 3 | 0.285116 (0.0921064,0.972282) |
| 4 | 0.000062 (0.000028,0.000266) |
| 5 | 0.00013862 (0.00002,∞) |
| 6 | 0.0073 (0.0047,0.0115) |
| 8 | 0.0044 (0.0011,0.025) |
| **9** | 0.0036 (0.002,0.0071) |
| 10 | 0.015 (0.0024,37) |
